# Antiviral activity of intracellular nanobodies targeting the influenza virus RNA-polymerase core

**DOI:** 10.1101/2023.08.31.555659

**Authors:** Mélissa Bessonne, Jessica Morel, Quentin Nevers, Allison Ballandras-Colas, Magali Grange, Alain Roussel, Thibaut Crépin, Bernard Delmas

**Affiliations:** Unité de Virologie et Immunologie moléculaires, INRAE, Université Paris-Saclay,Jouy-en-Josas, France; Institut de biologie structurale, CNRS, Université de Grenoble, Grenoble, France; Laboratoire d’Ingénierie des Systèmes Macromoléculaires (LISM), CNRS, Université d’Aix-Marseille, Marseille, France

**Author notes:** these authors contributed equally (Mélissa Bessonne, Jessica Morel).

## Abstract

Influenza viruses transcribe and replicate their genome in the nucleus of the infected cells, two functions that are supported by the viral RNA-dependent RNA-polymerase (FluPol). FluPol displays structural flexibility related to distinct functional states, from an inactive form to conformations competent for replication and transcription. FluPol machinery is constituted by a structurally-invariant core comprising the PB1 subunit stabilized with PA and PB2 domains, whereas the PA endonuclease and PB2 C-domains can pack in different configurations around the core. To get insights into the functioning of FluPol, we selected single-domain nanobodies (VHHs) specific of the influenza A FluPol core. When expressed intracellularly, several of them exhibited inhibitory activity on type A FluPol, but not on the type B one. The most potent VHH (VHH16) targets PA, but preferentially bind the PA-PB1 dimer with an affinity below the nanomolar range. Ectopic intracellular expression of VHH16 in virus permissive cells blocks multiplication of different influenza A subtypes, even when induced at late times post-infection. VHH16 was found to impair the transport of the PA-PB1 dimer to the nucleus, without affecting its handling by the importin β RanBP5 and subsequent steps in FluPol assembly. These data suggest that the VHH16 neutralization activity is likely due to an alteration of the import of the PA-PB1 dimer into the nucleus, resulting to an inhibition of FluPol functioning. VHH16 binding site represent a potential target for antiviral development.

**Author Summary:** The influenza virus RNA-polymerase (FluPol) ensures genome transcription and replication in the nucleus of the infected cells. To select ligands able to block FluPol activities, we screened a library of phages encoding nanobodies and resulting from the immunization of a llama with FluPol subunits. When expressed intracellularly, one of the nanobodies displays highly efficient FluPol blocking and virus neutralizing activities. This nanobody binds FluPol with high affinity and recognizes preferentially the PA-PB1 assembled subunits. Furthermore, it was found to interfere with the transport of the PA-PB1 dimer into the nucleus, suggesting that targeting FluPol trafficking between the cytoplasm and the nucleus may constitute a powerful strategy to develop new antivirals.

## Introduction

Seasonal human influenza viruses epidemics and pandemics in a recurrent mode cause significant morbidity and represent a main public health burden every year in the world. Furthermore, animal influenza viruses infecting pigs and avian species can compromise meat industries, as exemplified by the 50 million birds culled in the affected establishments with circulation of highly pathogenic avian influenza viruses in poultry, captive and wild birds in the 2021-2022 epidemic season in Europe [**1**].

Influenza A viruses genome is made of eight RNA segments of negative polarity packaged in viral ribonucleoprotein (vRNP) complexes (reviewed in [**2**]). Each of these vRNPs is composed of a large number of copies of the nucleoprotein (N) associated to genomic RNAs, with a single virus RNA-dependent RNA-polymerase (FluPol) bound to the 5’- and 3’-ends of the RNA segment. The vRNPs constitute the structural and functional units for genome replication and transcription which occur in the nucleus of the infected cell. The three largest segments encode the FluPol subunits: the two basic proteins PB1 and PB2 and the acidic subunit PA (reviewed in [**3**]). The polymerase subunits, which are produced in the cytoplasm, are then imported into the nucleus and assembled into a functional trimer [**4**]. PA and PB1 form a dimer in the cytoplasm, which is imported into the nucleus separately from PB2 [**5**], [**6**], [**7**], [**8**]. Once in the nucleus, the PB1-PA dimer associates with PB2 to form the heterotrimeric polymerase. The nucleotide polymerization activity is common to both replication and transcription, with an additional cap-snatching function being employed during transcription to steal short 5’-capped RNA primers from host mRNAs [**9**].

The PB1 subunit functions as the polymerase catalytic subunit. It binds to the promoter sequences of the viral and complementary RNAs, and catalyzes RNA chain elongation [**10**], [**11**]. The PB2 subunit is responsible for recognition and binding of the cap structure of host mRNAs [**12**], [**13**]. The PA subunit is divided into two main domains structurally well defined, the endonuclease domain (amino acids 1 to 197) and a large C-terminal domain (amino acids 257 to 716) that defines, with the PB1 subunit, the core of FluPol with the 237 N-ter residues of PB2. The PA endonuclease and the PB2 cap-binding domains are flexibly-linked domains to the core and they act synergistically to promote cap-snatching-dependent transcription [**14**].

Nanobodies (also known as VHHs), which are variable domains of heavy chain-only camelid antibodies of about 15kDa, have proved to be versatile molecular binders for their use as research tools in structural and cell biology [**15**], [**16**]. Their ability to bind their cognate ligand is not dependent of post-translational modifications such as disulfide bonds and glycosylation [**17**]. These properties allow the VHHs to be expressed in the cytosol of eukaryotic cells with preservation of their binding activity properties [**18**]. The intracellular expression of a VHH specific of a viral protein involved in any key function in the virus cycle may perturbate the target protein functioning and block virus multiplication (reviewed in [**19**]). Understanding the molecular bases involved in the inhibition of the virus replication may provide new perspectives in the design of antivirals targeting influenza viruses.

In this work, we generated nanobodies specific to the structurally rigid core domain of FluPol and identified a VHH able to block efficiently FluPol activity and virus multiplication (see **Fig. 1** for the rationale of the study). This VHH recognizes more efficiently the PA-PB1 dimer than its monomeric constituents, and interferes with the transport of PA-PB1 to the nucleus, without affecting the binding of RanBP5 to its dimeric target.

**Figure 1:**
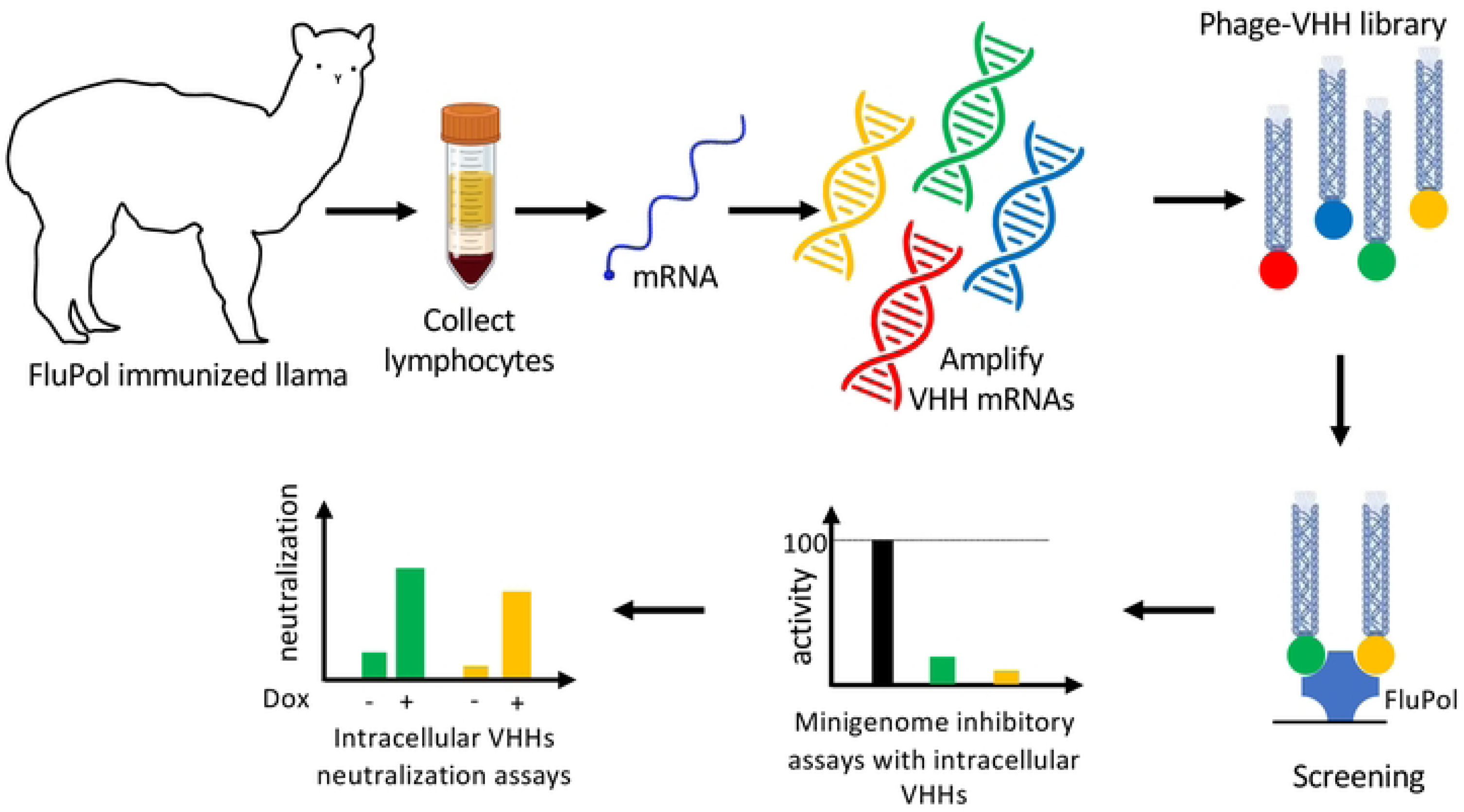
Selection mode of intracellular neutralizing FluPol-specific nanobodies/VHHs.

## Results

### Generation of VHHs specific to the influenza virus RNA-polymerase

To generate VHHs specific to the influenza virus RNA-polymerase, a llama was immunized with the core of WSN(H1N1) FluPol constituted by the full-length PB1 subunit with the two-third C-terminal moieties of PA (197-716) and PB2 amino acids (1-116) [**20**] (**Fig. 2A**). After five rounds of immunization, peripheral mononuclear cells were collected. VHH-encoding sequences were amplified from purified RNA and cloned into an M13 phagemid vector to create a specific nanobody phage library. Screening of nanobodies was performed by phage display as previously published [**15**], [**21**]. Three rounds of panning allowed the selection of seven FluPol specific binders (**Fig. 2B**).

**Figure 2:**
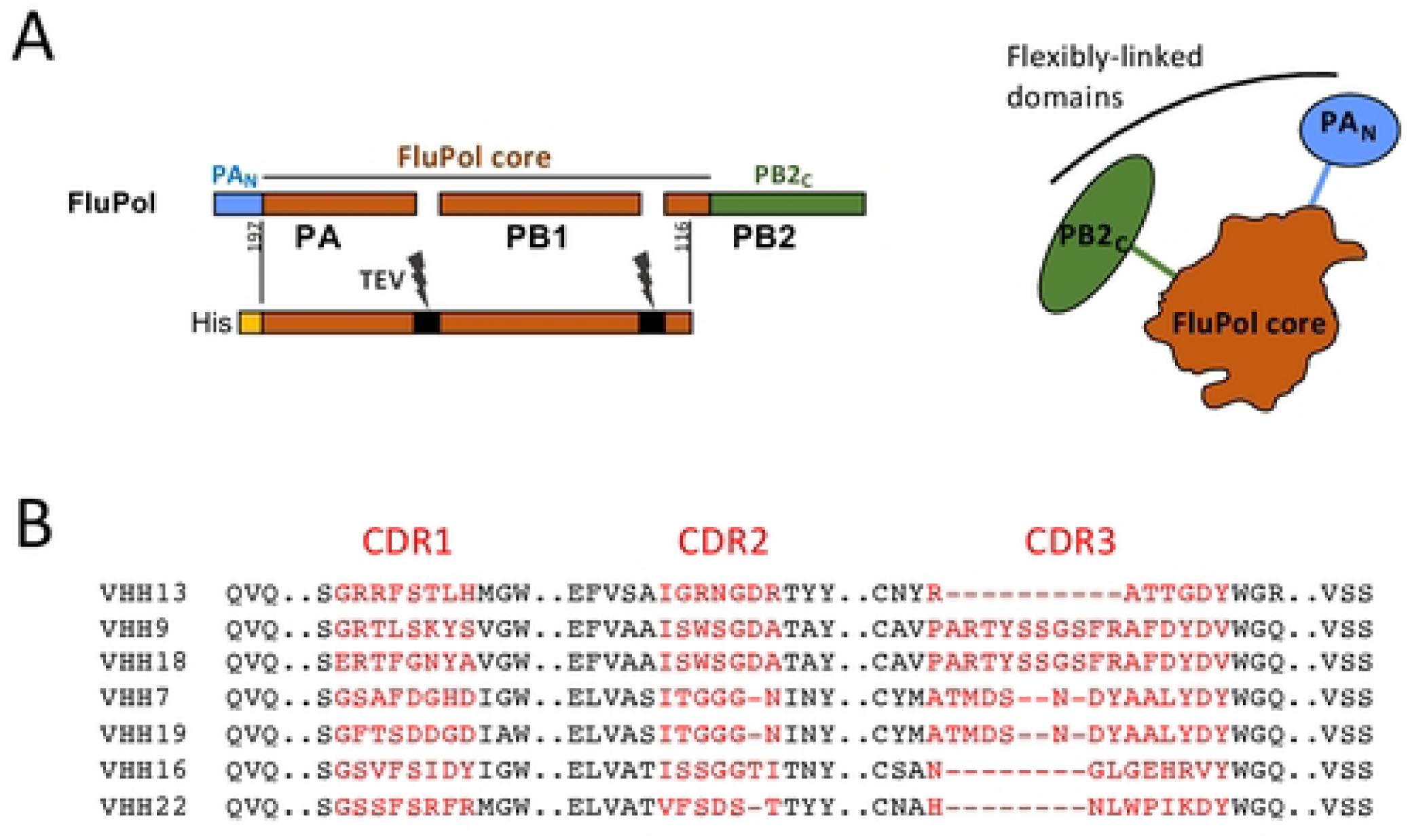
FluPol polypeptides and FluPol-specific VHH sequences. **A.** Schematic representation of FluPol and its structural domains, the core (colored in brown) constituted by PB1 and domains of the PA and PB2 subunits and flexibly-linked domains, PA_N_ and PB2_C_ (in blue and green colors) that display different packing arrangements onto the core. To select core-specific VHHs, the FluPol core was expressed as a fusion protein with a His-tag and linkers (black colors) cleavable by the TEV protease expressed in frame [**20**]. The core was produced and purified for alpaca immunization to generate a VHH library screening. **B.** ClustalW-based alignment of the anti-FluPol core VHHs with their CDR1, CDR2 and CDR3 domains. Strictly conserved residues are white on a red background, and partially conserved residues are red.

### Intracellular VHHs potency on FluPol activity

Using a minireplicon assay, we assessed the effects of intracellular expression of VHH cDNAs on the transcription/replication activity of the viral polymerase. The pPol1-WSN-NA-firefly luciferase plasmid produces a modified influenza NA viral RNA (vRNA) in which the NA-coding sequences are replaced by the firefly luciferase gene. Upon co-transfection of HEK-293T cells of the pPOL1-WSN-NA-firefly luciferase plasmid with plasmids allowing expression of PA, PB1, PB2 and NP with one VHH, the luciferase enzymatic activity measured in cell extracts reflects the overall transcription and replication activities of the transiently expressed viral polymerase. This assay revealed that two VHHs, VHH16 and VHH18 have strong potency in a cellular context, VHH16 being able to block up to 95% of FluPol activity of the H1N1 prototype WSN and an H3N2 virus (**Fig. 3A and S1 Fig**). In contrast, VHH16 did not block influenza B FluPol (**Fig. 3B**). We further established that FluPol inhibition by VHH16 was dose-dependent and that a 1/25 ratio between the plasmid expressing VHH16 and the ones expressing each FluPol subunit blocks 50% of the FluPol activity (**Fig. 3C**).

**Figure 3:**
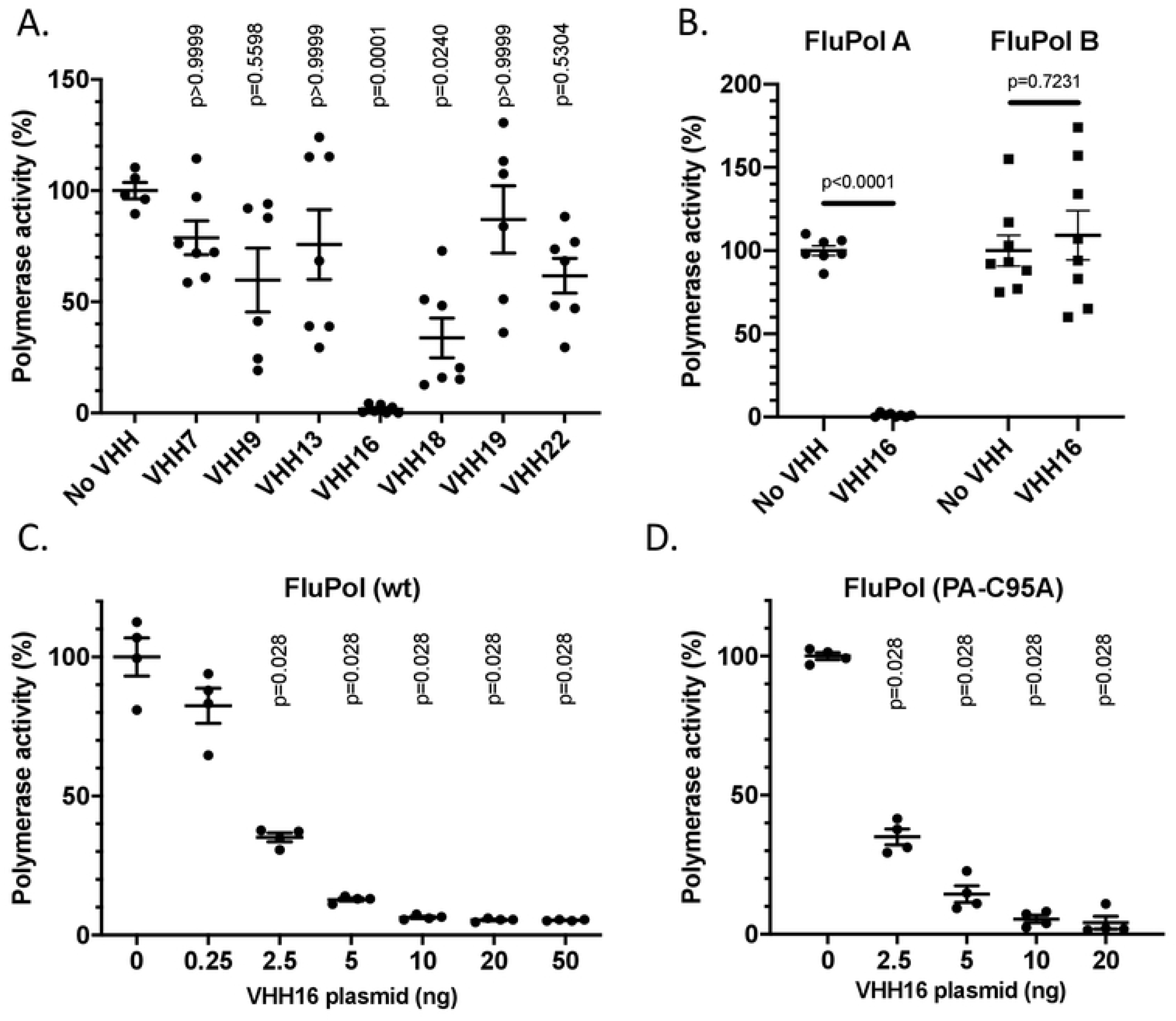
Effect of VHH expression on the FluPol replication and/or transcription activities. **(A)** Effect of VHHs on FluPol activity using a luciferase-reporter minireplicon assay. Plasmids expressing NP, PA, PB1, PB2 of the WSN strain (an influenza A H1N1 virus FluPol) were co-transfected in HEK 293T cells together with the WSN-NA-firefly-luciferase reporter plasmid and a plasmid encoding a VHH. A plasmid encoding the nano-luciferase was co-transfected to control DNA uptake and normalize minireplicon activity. Luciferase activities were measured in cell lysates 48 hours post-transfection. **(B)** Same procedure than in **(A)**, except that plasmids encoding the replicative complex of an influenza B virus (strain B/Memphis/13/2003) with its replicon was included in the assay. In (**A and B**), data are mean ± s.e.m. n=2 independent transfections with n=3 or 4 technical replicates. Kruskal-Wallis test was used to compare luminescence in the presence or absence of nanobodies. p<0.05 is considered significant. In (**C**), the FluPol activity was quantified in transfections containing 0 to 50 ng of the VHH16 plasmid per P96 well. An additional empty plasmid was included in each transfection to transfect the same amount of DNA. Data are mean ± s.e.m. with n=4 technical replicates. In panel (**D**), the same procedure than in (**C**) was used, except that a plasmid encoding the PA C95A mutant deficient in replicase activity was used instead of the one encoding the PA wild-type subunit. Data are mean ± s.e.m. with n=4 technical replicates. Kruskal-Wallis test was used to compare luminescence in the presence or absence of nanobodies. **(C and D)** Kruskal-Wallis test was used to compare luminescence in the presence or absence of VHHs. p<0.05 is considered significant.

### VHH16 inhibits influenza A virus multiplication *in cellulo*

Having shown the ability of several VHHs to inhibit FluPol activity in cells, we wondered whether they would be able to inhibit viral multiplication. For this purpose, we used a recombinant bioluminescent reporter virus, with the nanoluciferase gene inserted in frame into the PA genomic segment, to quantify viral multiplication [**22**]. We first assessed the inhibitory effect of VHHs in a transient transfection assay in HEK-293T cells, in which VHH-encoding plasmids were transfected 24 hrs before infection (**S2 Fig**). While VHH18, 7 and 9 were not able to block virus multiplication in this assay, VHH16 displays an inhibition activity on FluPol as evidenced by a lower activity of the luciferase reporter in infected cells.

To confirm this result, we aimed at determining the virus susceptibility of MDCK cell lineages constitutively expressing VHH16. Thus, a construct in which the VHH16 open reading frame was fused to sequences encoding a 2A self-cleaving peptide and GFP was placed under the control of a constitutive promoter in a lentiviral vector. MDCK cells were transduced with the resulting plasmid or with a lentivirus driving expression of GFP alone. **S3 Fig** exemplifies the stable expression of the VHH16-2A-GFP construct in selected MDCK clones. Infections of serial dilutions of the virus inoculum revealed a marked block of infection in MDCK-VHH16 cells when compared to the MDCK-GFP cells (**Fig. 4A**). A 10^4^-fold decrease of virus multiplication was evidenced at 48 hrs post infection with a multiplicity of infection of 0.005. The cytopathic effect of the infection was found delayed or even blocked in MDCK-VHH16 cell monolayers, compared to MDCK-GFP cells (**Fig. 4B**).

**Figure 4:**
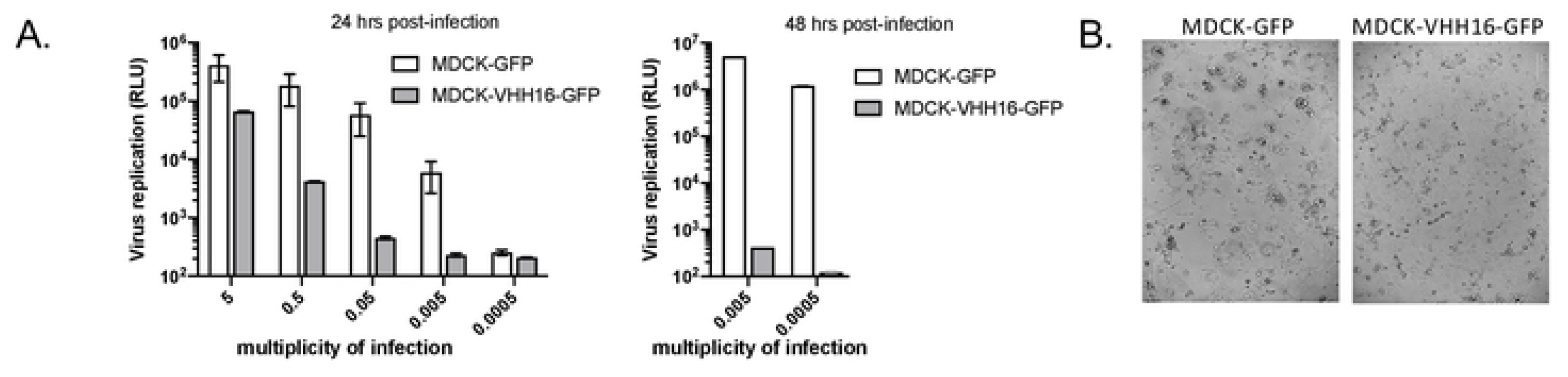
MDCK cells constitutively expressing a GFP-tagged version of VHH16 (or GFP as control cells) were seeded 24 h before infection with an influenza A virus encoding a reporter nanoluciferase (WSN-Luc). **(A)** Infections were carried out at different multiplicities of infection (m.o.i.) and virus replication was quantified in cell lysates 24 hours and 48 hours post-infection. **(B)** Light microscopy views of MDCK-VHH16-GFP and MDCK-GFP cells infected at a m.o.i. of 0.05 24 hours post-infection.

To analyze more finely the ability of VHH16 to block influenza virus multiplication, we engineered several additional cell lines, in which the VHH16-2A-GFP construct was placed under the control of the P_TGE3G_ promoter, which is inducible by doxycycline (**Fig. 5A**). Thus, with these constructs, we may be able to quantify the neutralization activity of VHH16 when expressed at early or late stages of infection. As exemplified in **S4 Fig**, expression of VHH16 was revealed only under doxycycline expression induction in RK13, a rabbit influenza virus permissive cell line. To probe the effect of VHH16 expression on infection, doxycycline was added 24 hrs before infection in cultured RK13-VHH16 and MDCK-VHH16 clones. While virus multiplication was effective in uninduced cells, Dox-induced RK13 and MDCK cell clones expressing VHH16 were found refractory to influenza virus infection (**Fig. 5B and S5 Fig**). A dose-response quantification of virus replication as a function of doxycycline concentration showed that 0.04 μM of doxycycline blocked efficiently virus multiplication (**Fig. 5C**). Next, we determined the effects of VHH16 expression when induction was carried out at different times post-infection (**Fig. 5D**). Strikingly, doxycycline addition at 6 hrs post-infection results in a block of 50 % of virus replication.

**Figure 5:**
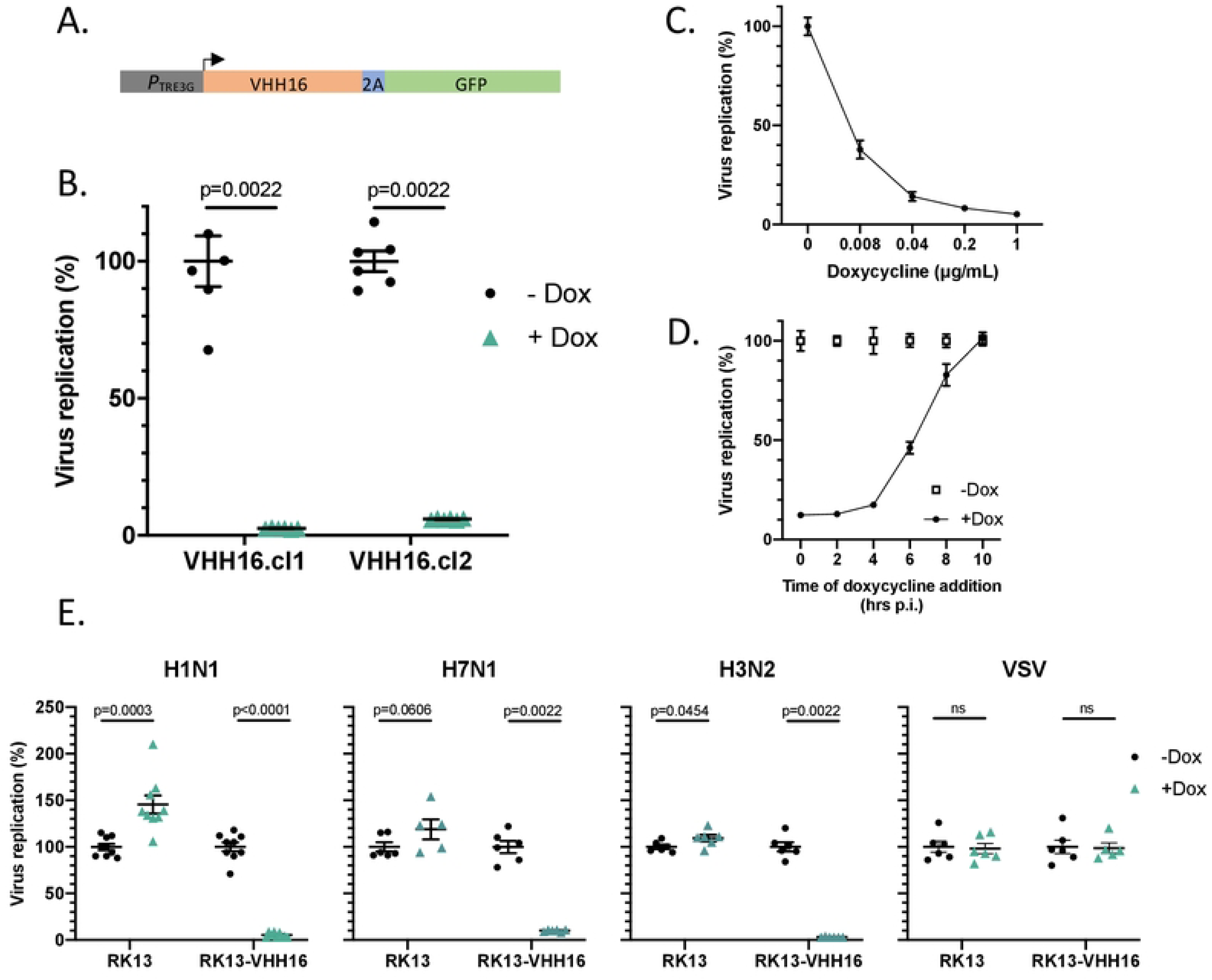
(**A)** Scheme of the DNA construct to promote VHH16-2A-GFP gene expression in a doxycycline (Dox)-inducible manner. (**B)** Two different RK13 cell clones selected for VHH16-2A-GFP gene expression were incubated (or not) with doxycycline and infected with the reporter influenza virus WSN-Luc. Twenty-four hours post-infection, virus replication was quantified by measurement of the luciferase activity. Data are mean ± s.e.m. n=2 independent transfections with n=3 technical replicates. 2-way ANOVA test was used to compare luminescence in the presence or absence of doxycycline. p<0.05 is considered significant. **(C)** WSN-Luc virus replication quantification as a function of Dox concentration. **(D)** Quantification of WSN-Luc virus replication when Dox (1 μg/mL) was added at different times post-infection. **(E)** Replication quantification of influenza virus types (H1N1, H7N1 and H3N2) encoding a reporter nanoluciferase in RK13 and RK13-VHH16 cells incubated with doxycycline. VSV-dsRed virus [**39**] replication was quantified by measuring mCherry fluorescence in fixed cells using a TECAN spectrophotometer Infinite 200 PRO (excitation wave length 580 nm for an emission wave length lecture at 620 nm). VHH16 expression did not inhibit VSV-dsRed replication.

To further investigate the spectrum activity of VHH16, we determined its neutralization activity towards two additional influenza viruses belonging to the H7N1 and H3N2 subtypes (A/Turkey/Italy/977/1999 [H7N1] and human influenza A/Scotland/20/1974 [H3N2]), and both tagged with the nanoluciferase (Nluc) reporter gene (**Fig. 5E**). As found with the H1N1 virus (WSN strain), multiplication of these two viruses was blocked efficiently when VHH16 expression was induced, suggesting a sequence conservation of the VHH16 epitope. Replication of vesicular stomatitis virus (a virus unrelated to influenza viruses) was not altered after doxycycline induction.

### VHH16 interacts specifically with the PA-PB1 dimer with high affinity

To identify the FluPol subunit recognized by VHH16, we used the previously described *Gaussia princeps* luciferase-based complementation assay [**23**]. In this assay, an interaction between two proteins each fused to either the Luc1 or the Luc2 segments of the *Gaussia* luciferase enzyme, results in reconstitution of a functional luciferase activity, which can be quantified by addition of the substrate. To favor proper folding of the FluPol subunits and the VHH16, the two luciferase moieties were fused onto their C terminus. For each sample, the specificity of the interaction over background was estimated by calculating a normalized luminescent ratio (NLR, [**24**]). Co-expressed FluPol subunits PA, PB1 or PB2 (fused to Luc1) with VHH16 (fused to Luc2) revealed a NLR value of 10 for the PA-VHH16 interaction (**Fig. 6A**) and no interaction between PB1 or PB2 with the VHH16. When PA-Luc1 was co-expressed with PB1 and VHH16-Luc2, NLR value reached 175, suggesting that VHH16 recognizes preferentially the PA-PB1 dimer than the monomeric PA. The interaction between VHH16 and the PA-PB1 dimer was also revealed when PB1 fused to Luc1 was co-expressed with PA. We next explore the VHH16 affinity for the PA-PB1 dimer by biolayer interferometry (BLI) at different concentrations to determine its kinetic rate constants (**Fig. 6B**). VHH16 displays low on- and off-rate constants, resulting in a dissociation constant that cannot be precisely estimated, but below the nanomolar range and corresponding to an almost non-reversible interaction.

**Figure 6:**
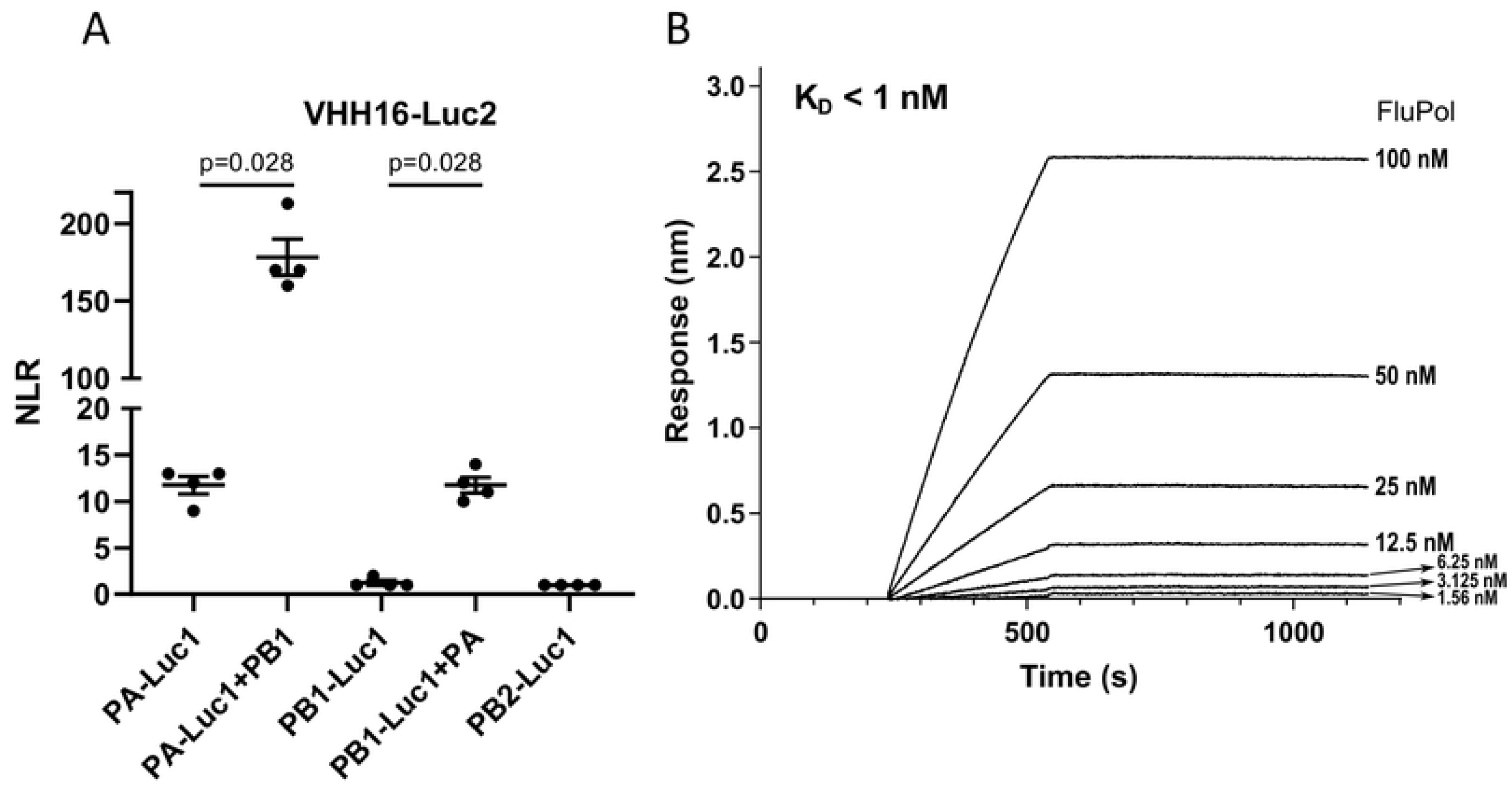
The interaction between the VHH16 and FluPol. **(A)** Complementation split-luciferase interaction assay between VHH16 and FluPol subunits. The normalized luminescence ratio (NLR) is calculated as described in the Materials and Methods section to quantify the interaction between VHH16 fused to Luc1 and FluPol subunits fused to Luc2. Luc1- and Luc-2 tagged polypeptides were expressed with or without untagged FluPol subunits. Twenty-four hours post-transfection, cells were lysed and luminescence was measured. Data are mean ± s.e.m. n=4 technical replicates and are representative of several experiments. Mann-Withney test was used to compare the NLR values, p<0.05 is considered significant. **(B)** BLI binding kinetics measurements between VHH16 and the PA-PB1 dimer. Equilibrium dissociation constants (K_D_) was determined on the basis of fits, applying a 1:1 interaction model.

### VHH16 inhibits FluPol transcription

As shown above, VHH16 inhibits the transcription/replication activity of FluPol when expressed without other viral components (**Fig. 3**). We next confirmed in an infection assay that transcription and replication were impaired in cells constitutively expressing VHH16. Viral RNA quantification showed a decrease of mRNA, cRNA, and vRNA levels when compared to control cells (**S6 Fig**). These results suggest that VHH16 neutralization activity may be due to a block of its RNA-polymerase activity in the nucleus, targeting transcription and/or replication, or any other earlier steps in the virus cycle such as FluPol assembly or the import of its subunits in the nucleus. Next, to determine if VHH16 neutralization activity inhibits transcription, we first assessed the transcriptional-only activity of FluPol when VHH16 was co-expressed. For this purpose, we took advantage of the previous identification of a FluPol mutant (PA-C95A) that is defective for replication [**25**]. Using the minireplicon assay used above with this FluPol PA-C95A mutant, we evidenced that FluPol transcription is inhibited by VHH16 in a dose-dependent manner (**Fig. 3D**) with the same magnitude than in the assay that integrate replication + transcription activities of FluPol (**Fig. 3C**), suggesting no specificity between replication and transcription in its mode of action.

### VHH16 does not block PA-PB1 dimer / PB2 assembly

Following their synthesis in the cytoplasm, while PA and PB1 are transported in the nucleus as a dimer, PB2 is imported in this compartment separately [**5**], [**6**]. Once in the nucleus, the PA-PB1 dimer associates with PB2 to form the functional heterotrimeric polymerase. To determine whether VHH16 binding on the PA-PB1 dimer may interfere with its association of PB2 in the nucleus, we used a protein complementation assay between the PA-PB1 dimer and PB2. As shown in **Fig. 7A**, while PA alone does not bind to PB2 in the absence of PB1, it associates with in the presence of PB1 (with an NLR value of 100). Co-expression of VHH16 or VHH7 with the three FluPol subunits did not modulate final FluPol assembly.

**Figure 7:**
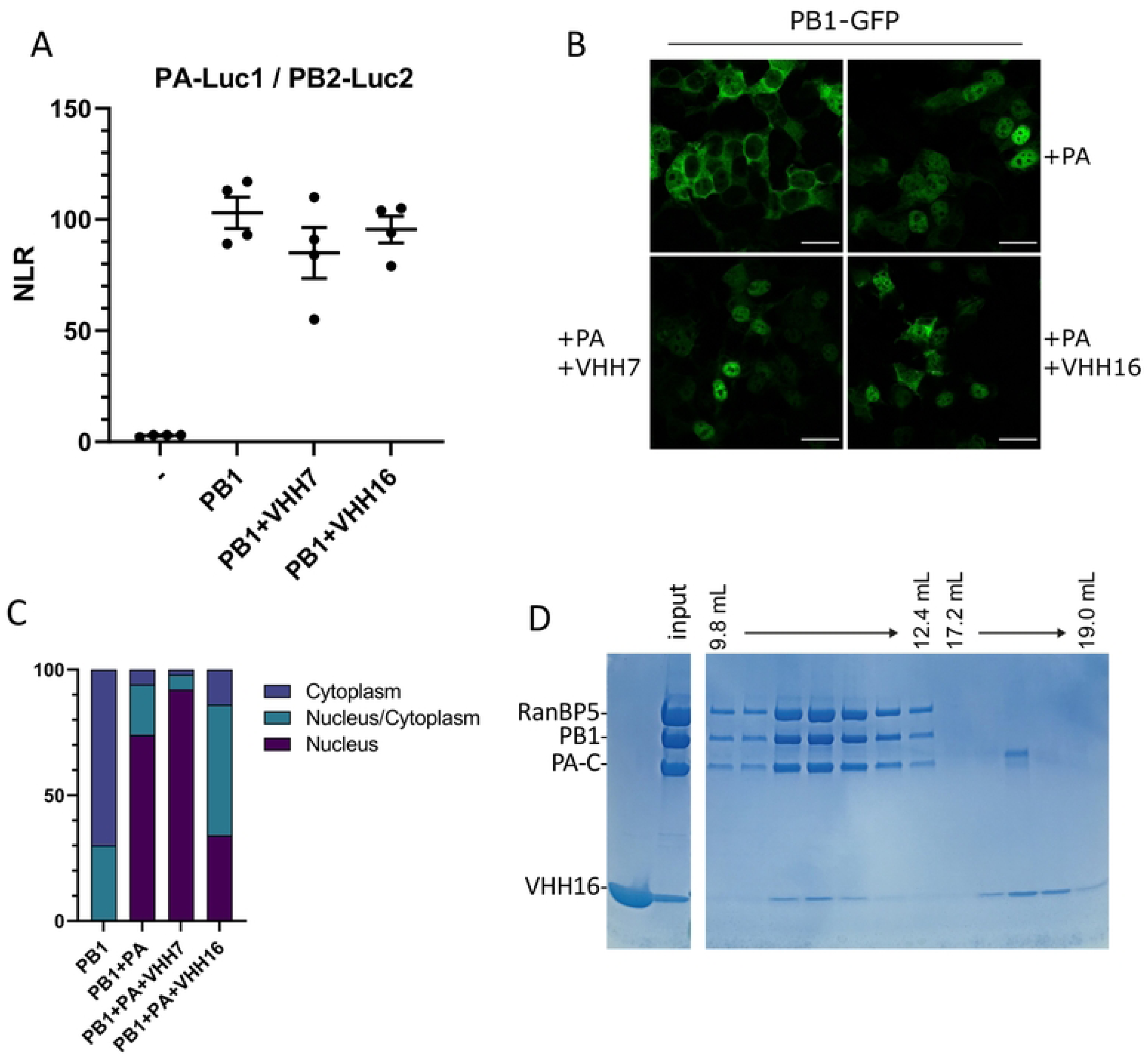
FluPol trafficking and assembly in the presence of VHHs. **(A)** Complementation split-luciferase interaction assay between PA-Luc1 and PB2-Luc2 fusion proteins in the presence of PB1 and VHHs as indicated. The normalized luminescence ratio (NLR) is calculated as described in the Materials and Methods section. Twenty-four hours post-transfection, cells were lysed and luminescence was measured. Data are mean ± s.e.m. n=4 technical replicates and are representative of several experiments. **(B)** Subcellular localization of the PB1-GFP fusion protein co-expressed with PA and VHHs. **(C)** Percentage of PB1-GFP-expressing cells with nuclear versus cytoplasmic localization of PB1-GFP. Fifty GFP-positive cells were scored for each condition. **(D)** Evidence for a complex constituted by the PB1 subunit associated to PA_linker+C_, RanBP5 and VHH16. A PA_linker+C_-PB1-RanBP5 complex was produced in insect cells and purified before incubation with excess of VHH16. Gel filtration was carried out and aliquotes of fractions were analyzed by SDS-PAGE and Coomassie blue staining. VHH16 was found associated to FluPol-RanBP5 complexes.

### VHH16 interferes with the transport of the PA-PB1 dimer to the nucleus

The inhibition of the RNA-polymerase activity of FluPol by VHH16 could result from a block in the genome transcription/replication process or in any upstream step, such as the assembly of the polymerase subunits or the import of the polymerase subunits in the nucleus. In order to determine whether VHH16 may interfere with the nuclear import of the PA-PB1 dimer into the nucleus, we used plasmids to co-express a fluorescently labeled PB1 (PB1-GFP, [**23**]) and PA with VHH16 to visualize the localization of the PB1 subunit. Our experiments confirmed that when PA and PB1 were expressed together, PB1 localized into the nucleus, in contrast to what observed when PB1 was expressed alone (**Fig. 7B and 7C**). Seventy-three percent of the labeled cells display an exclusive nucleus labeling when PA and PB1 were co-expressed. When PB1-GFP and PA were co-expressed with VHH16, only thirty-two percent of the cells exhibited an exclusive fluorescence signal in the nucleus. When PB1 and PA were co-expressed with VHH7 (a non-neutralizing VHH), 90% of the cells display a strict nucleus labeling. We concluded from these data that VHH16 binding interfere with the import of the PA-PB1 dimer in the nucleus. We next hypothesized that VHH16 may interfere with the binding of RanBP5, the β-importin that supports the transport of the PA-PB1 dimer to the nucleus [**26**].

### VHH16 does not displace the interaction of the importin RanBP5 with the PA-PB1 dimer

To determine if VHH16 may displace RanBP5 complexed with the PA-PB1 dimer, a recombinant equimolar complex made by PA and PB1 subunits with RanBP5 was purified as described previously [**20**] and incubated with the VHH16 to analyze its impact on the complex. **Fig. 7D** shows that VHH16 was able to bind efficiently to the PA-PB1-RanBP5 complex, but did not displace RanBP5 from the PA-PB1 dimer. This result shows that VHH16 does not interfere with RanBP5 binding and the VHH16 binding site does not overlap the one of RanBP5 on PA-PB1.

## Discussion

In this study, we exploited the unique properties of single-domain antibodies to identify new vulnerabilities in the functioning of FluPol. While their biosynthesis includes secretory pathway post-translational modifications, such as glycosylation and disulfide bond formation, their ability to recognize an epitope is generally well conserved when they are expressed in the cytosol of eukaryotic cells or in bacteria. We thus generated VHHs targeting FluPol and studied their impact on its activity when they were expressed in the cytosol of infected cells. While three of them were able to inhibit transcriptional/replicase activity, only VHH16 was able to block virus multiplication in a transient expression assay. Its neutralization activity was found highly potent in cell clones constitutively expressing the VHH. Moreover, VHH16 expression induction at late time of infection still resulted in a block of infection, demonstrating its potentialities. Its affinity (that was impossible to quantify precisely in the picomolar range due to a too low non-measurable off-rate constant) may account for its high neutralization activity. Another striking feature of the interaction of VHH16 with FluPol in a cellular context is its more efficient binding on the PA+PB1 dimer than to PA alone, suggesting that PB1 may stabilize PA structure, and as a consequence, VHH16 binding site on PA. Thus, the PA linker, a flexible domain that is located between the endonuclease and the PA-C domain and that is structured when PA is associated to PB1, may constitute the VHH16 epitope. It is interesting to note here that an anti-PA Mab was previously found to exhibit an enhanced reactivity to the PA-PB1 complex, also suggesting that PA-PB1 interactions induce a conformation change in PA, which could be required for its nuclear translocation [**8**]. Another possibility is that some PB1 residues may also contribute to VHH16 epitope at a PA-PB1 junction. Such overlap of VHH epitopes on a PA-PB1 junction has been previously recognized with other anti-FluPol inhibitory nanobodies [**27**].

How does VHH16 inhibit virus multiplication? Several hypotheses have been investigated to identify the function(s) of FluPol that are blocked or inhibited by VHH16. Our data show that VHH16 does not inhibit the association of PB2 to the PA-PB1 dimer, a step in virus assembly that occurs in the nucleus. Our results suggest that the inhibitory activity of VHH16 is related to a perturbation of the import of the PA-PB1 dimer into the nucleus, this activity being not associated to a block of the binding of the importin β RanBP5 to the dimer. At the same way, it is interesting to note that a VHH targeting the NP of influenza A virus exhibit antiviral activity by blocking nuclear import of incoming viral ribonucleoproteins, and as a consequence, both viral transcription and replication in the nucleus [**28**], [**29**]. It should also be reminded that nucleozin, a potent inhibitor of influenza virus infection, triggers NP aggregation and antagonizes its nuclear accumulation [**30**]. Thus, the trafficking of FluPol and/or the replication complex between the cytosol and the nucleus can be considered for potential influenza antiviral development.

It is also interesting to note that the transport of the PA-PB1 dimer into the nucleus constitutes a critical step in temperature-sensitive influenza A virus mutants generated in the PA linker [**23**], [**31**]. In these mutants, while their production and the transport of their PA+PB1 subunits is not impaired at 33°C and 37°C, virus multiplication and the PA+PB1 transport into the nucleus is blocked at 39.5°C, thus confirming the vulnerability of influenza viruses at this step of the virus cycle.

However, our data do not rule out that VHH16 may also interfere with viral transcription and replication *per se*. For instance, FluPol require interactions with different host factors to ensure viral transcription and replication, such as RNA-polymerase II and ANP32A, respectively (see [**9**], [**32**], [**33**], [**34**] and references therein). VHH16 may interfere with the association of these factors with FluPol to inhibit these functions.

Since VHH16 is able to block the replication of several viruses representative of the H1N1, H3N2 and H7N1 subtypes, the VHH16 epitope may constitute an effective target for potential influenza antiviral therapy. Its structural characterization should be carried out to contribute to the design of ligands retaining the inhibitory activity of VHH16 and able to cross the cell membrane barrier.

## Materials and methods

### Cells

Human embryonic kidney HEK-293T cells (ATCC) were grown in complete Dulbecco’s modified Eagle’s medium (DMEM, Eurobio scientific) supplemented with 10% fetal calf serum (FCS, Capricorn) and 1% penicillin-streptomycin. Madin-Darby canine kidney (MDCK) and rabbit kidney (RK13) cells were grown in modified Eagle’s medium (MEM, Eurobio scientific) supplemented with 5% fetal calf serum, glutamine, and penicillin-streptomycin. For MDCK and RK13-derived clones, 4 or 1 μg/ml of puromycin was added to the cell culture medium, respectively.

### Plasmids

VHH ORFs were cloned in pCI mammalian expression vector (in Nhe1-Mlu1 restriction sites; Promega). The VHH16 ORF was also fused in frame with sequences encoding the 2A peptide of porcine teschovirus 1 and GFP and placed under the control of a constitutive promoter in a lentiviral vector – the 2A peptide allowing the co-translational cleavage between VHH16 and GFP. The VHH-2A-GFP ORF was also placed under the control of the P_TREG_ promoter, which is inducible by doxycycline (Takara Bio). For minigenome assays, a pPolI-Firefly plasmid encoding the Firefly luciferase sequence in negative polarity flanked by the 5’ and 3’ non-coding regions of the IAV NA segment was used [**23**]. The pCMV-Nanoluc (Promega) plasmid was used as an internal control to normalize transfection efficiency.

### MDCK- and RK13-VHH16 cell clones selection

Cell clones were either produced by lentiviral transduction or plasmid transfection. For lentiviral production, 293T cells (ATCC) were co-transfected with pLV-EF1a-IRES-puro (Addgene 85132) encoding GFP or the fusion protein VHH16-2A-GFP, packaging plasmid R8.2 (Addgene 12263) and VSV-G plasmid (Addgene 8454) as described [**35**]. Supernatants containing vectors were harvested at 36h, 48h and 72h, clarified and filtered. MDCK cells in 6-wells plate were transduced for 24h with lentiviral vectors and puromycin selection was applied 48h after transduction to generate MDCK-VHH16 and MDCK-GFP cells.

MDCK and RK13 cells inducibly expressing HA-tagged VHH16 were generated by co-transfection of pCMV-TET3G, pTRE3G-VHH16 and a plasmid expressing the puromycin resistance cassette and selected in the presence of puromycin (1 μg/mL for RK13- and 4 μg/mL for MDCK-cell clones).

### Protein expression and purification

Expression and purification of the FluPol was done as previously described [**20**]. Briefly, large scale suspension cultures expressing the polymerase fusion constructs of A/Victoria/3/1975(H3N2) was prepared using High Five insect cells grown in Express Five media (Life Technologies) at 0.5 × 10^6^ cells/mL infected at 0.2% (V/V) with the baculovirus mother solution. Cultures were maintained at 0.5–1 × 10^6^ cells/mL until proliferation arrest (24–48 h after infection). Cultures were then spun down at 800 g for 10 min and cell pellets were stored at − 80 °C.

Cell pellets were resuspended in 50 mL of lysis buffer (50 mM Tris-HCl pH 8.5, 300 mM NaCl and 2 mM β -mercaptoethanol) per 5 × 10^8^ cells in the presence of EDTA-free anti-protease cocktail (complete from Roche). Lysis was performed with two cycles of freezing (− 180 °C) / thawing (26 °C) after which 10 % of glycerol were added to the lysate before centrifugation (45 min, 40 000 g, 4 °C). After retrieval of the clarified lysate, 15 mM of imidazole pH 8.0 were added before loading on Ni-NTa superpose resin (Quiagen). After flowing the lysate through the resin, wash steps were performed and bound complex was eluted with 300 mM imidazole. Elution fractions containing the polymerase core were pooled and dialysed overnight against a 20 mM Hepes pH 7.5, 300 Mm NaCl, 5 Mm betamercaptoethanol, 10% glycerol buffer before being subjected to a Hitrap heparin resin (Cytiva). After binding to the resin and wash steps, elution was carried out with a salt gradient on a FPLC system. The FluPol elution peak was pooled and flash freezing in liquid nitrogen.

### VHH library generation

Four injections of 0.5 mg purified FluPol core (in 20 mMTris–HClpH 8.0, 150 mMNaCl) were performed subcutaneously at one-week intervals followed by a fifth injection two weeks later in one llama (*Lama glama*; from Ardèche Lamas, France). Lymphocytes were isolated from blood samples obtained 5 days after the last immunization. The cDNA was synthesized from purified total RNA by reverse transcription and was used as a template for PCR amplification to amplify the sequences corresponding to the variable domains of the heavy-chain antibodies. PCR fragments were then cloned into the phagemid vector pHEN4 [**36**] to create a nanobody phage display library. Selection and screening of nanobodies were performed as previously described [**15**]. Three rounds of panning resulted in the isolation of FluPol-specific binders. VHH−7, −9, −13, −16, −18, −19 and −22 were selected and sequenced. The synthetic genes corresponding to the VHH sequences with a pelB signal peptide and a C-terminal 6xHis tag were obtained from Twist Bioscience and cloned in between *EcoRI* and *NotI* sites in a pET-28a expression vector.

### Protein complementation and minigenome assays

HEK-293T cells were seeded in 48 well plates (1×10^5^ cells/well) one day before being transfected with 150ng pci-VHH16-Luc2 together with 150ng of either pci-PA-Luc1, pci-PB1-Luc1 or pci-PB2-Luc1 and another 150ng of either pci-neo, pci-PB1 or pci-PA using polyethylenimine (PEI). 24h later, cells were lysed using Renilla lysis buffer (Promega) for 20min at room temperature under shaking (250rpm) and luminescence was quantified with the Renilla Luciferase Assay System (Promega).

HEK-293T cells were seeded in 96 well plates (5×10^4^ cells/well) one day before being transfected with 25 ng of each of the pCI plasmids expressing the PB2, PB1, PA viral proteins, together with 50, 10 and 5ng of the pCI-NP, pPolI-Firefly and pCMV-Nanoluc plasmids, respectively, and 10ng or a range from 0 to 50ng pci-VHH using PEI. 48h later, cells were lysed using PLB buffer (30 mM Tris pH7.9, 10 mM MgCl_2_, 1% Triton X-100, 20% Glycerol, 1 mM DTT) buffer and luminescence activity was measured with the Luciferase Assay System (Promega).

### Viruses

The recombinant reporter virus WSN(H1N1)-nanoluciferase (H1N1-Nluc, previously named PASTN or PA-SWAP-2A-NLuc50 in [**22**]) was kindly provided by Andrew Mehle. In this reporter virus, the “self-cleaving” 2A peptide from porcine teschovirus and the nanoluciferase coding sequence were placed downstream of the PA sequence to create a contiguous ORF. Native packaging sequences were restored by repeating the terminal 50 nucleotides of the PA ORF (including the stop codon) after the Nluc stop codon adjacent the native untranslated region. Direct repeats were removed from the reporter gene by introducing silent mutations at the 3’ end of the PA ORF. Reverse genetic systems for avian influenza A/Turkey/Italy/977/1999 [H7N1] and human influenza A/Scotland/20/1974 [H3N2] viruses have been previously elaborated and used to generate the recombinant reporter viruses H7N1-Nluc and H3N2-Nluc [**37**], [**38**]. Both these Nluc reporter viruses were designed using the same strategy than H1N1-Nluc, with their PA segments encoding a PA-2A-Nluc polyprotein, and were kindly provided by Ronan Le Goffic. The reporter VSV-mCherry virus [**39**] in which the fluorescent protein mCherry ORF was inserted in the L-protein reading frame was kindly provided by Emmanuel Heilmann.

### Infection assays

#### in cells transiently expressing VHHs

HEK-293T cells were seeded in a 6-well plates at a density of 0.8 x 10^6^ cells. Cells were transfected with 2,5μg of pCI plasmid encoding VHH7, VHH9, VHH16, VHH18 or an empty pCI plasmid, using PEI. Subsequently, 20h post-transfection, cells were infected with a WSN-PA-nanoluc virus at a multiplicity of infection of 0,01. Cells were lysed 24h after, using PLB buffer and luminescence activity was measured with the Nano-Glo® Luciferase Assay System (Promega).

#### in MDCK and RK13 cell clones

Cells were seeded in 12-well plates at a density of 0.3 x 10^6^. In (Dox+)-inducible cells, doxycycline was added at 1μg/mL (or at the indicated concentration) 24h before infection to induce VHH expression. Cells were infected with influenza viruses tagged with a reporter gene (nanoluciferase) or by VSV-mCherry at a multiplicity of infection of 0.001. Virus replication was quantified by measuring nanoluciferase luminescence and mCherry fluorescence activities.

### Immunofluorescence assay

HEK 293T cells were seeded in 24-well plates on coverslips 24 hours before being transfected with 600ng pci-PB1-GFP, pci-PA and pci-VHH16 total. After 24 hours, cells were fixed with 4% paraformaldehyde (PFA) and VHH16 was revealed using a secondary antibody coupled to AlexaFluor 546. Coverslips were then visualized by confocal microscopy.

### Affinity determination by Bio layer interferometry

Binding kinetics experiments were performed on an Octet system (Octet RED96) (FortéBio, CA). A black bottom 96-well microplate was filled with 200 μL of solution (PA-PB1 dimer in PBS buffer) and agitated at 1,000 rpm, and all experiments were carried out at 25°C. Tips were hydrated in PBS buffer for 1 hour at room temperature prior experiments. Biotinylated VHH16 (4 μg/mL) were loaded on streptavidin SA (18–0009) biosensors (Pall ForteBio) for 1 min. After a baseline step in assay buffer (PBS [pH 7.4], 0.1% bovine serum albumin, 0.02% Tween 20), ligand-loaded sensors were dipped into known concentrations of PA-PB1 dimer for an association phase during 500 to 700 seconds. The sensors were then dipped in assay buffer for a dissociation step during 1000 seconds in assay buffer. Association and dissociation curves were globally fitted to a 1:1 binding model. Binding curves were fit using the “association then dissociation” equation in the FortéBio Data analysis software version 7.1 to estimate the K_D_.

### Viral RNA quantification by RT-PCR

Strand-specific real-time RT-PCR was carried out as previously described using primers specific of segment 5 [**40**]. The primers used are listed in Table 1. Briefly, cDNAs were synthesized with strand-specific RT primers tagged at their 5′ end with the hot-start modification of using saturated trehalose: a 5.5 μL mixture containing 200 ng of total RNA sample and 10 pmol of tagged RT primer was heated for 10 min at 65°C, chilled immediately at 4°C for 5 min, and then heated again at 60°C. After 5 min, 14.5 μL of a preheated reaction mix (4 μl First Strand buffer 5X (Thermo Fisher), 1 μL 0.1 M dithiothreitol (DTT), 1μL dNTP mix (10 mM each), 1μL Superscript III reverse transcriptase (200 U/μl, Thermo Fisher), 1 μL RNasin Plus RNase inhibitor (40 U/μL, Promega) and 6.5 μL saturated trehalose) was added and incubated at 60°C for 1 h. Real-time PCR (qPCR) was performed with the iTaq™ Universal SYBR® Green Supermix (BioRad) on a CFX96 (BioRad). Seven microliters of a 10-fold dilution of the cDNA was added to the qPCR reaction mixture (10 μL Brilliant II SYBR Green qPCR Master Mix 2X, 1.5 μL of each qPCR primer at 10 μM). The cycle conditions of qPCR were 95°C for 5 min, followed by 50 cycles of 95°C for 15 s, 56°C for 15 s and 60°C for 45 s.

**Table 1:**
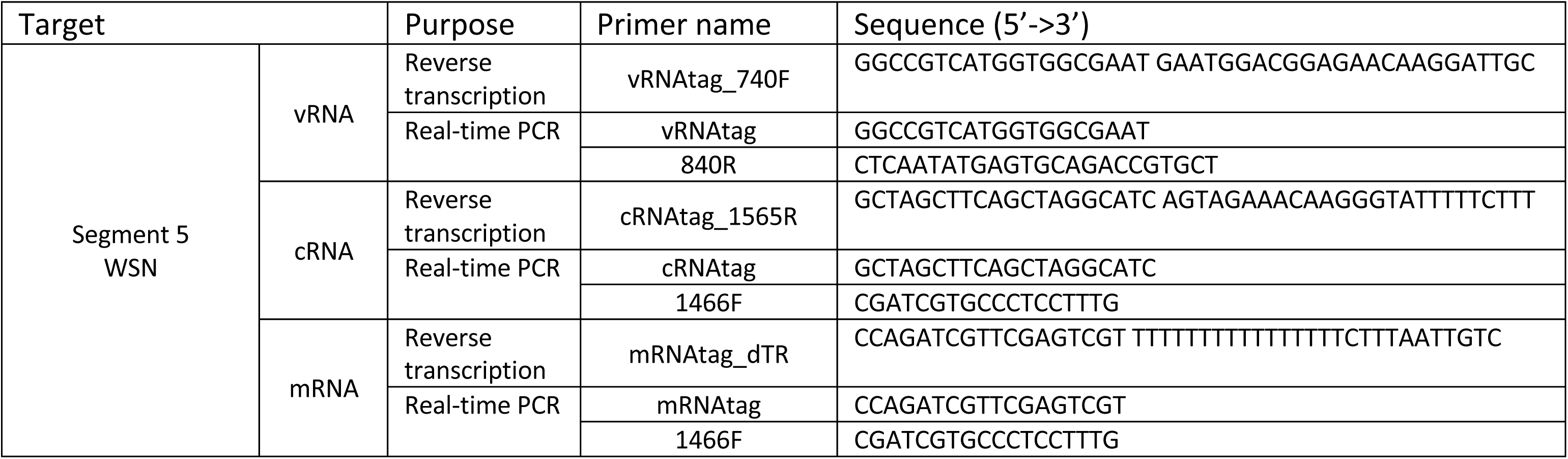
Primer sets for strand-specific real-time RT-PCR.

## Acknowledgments

We thank Andrew Mehle for the gift of the reverse genetics system of the IAV H1N1-Nluc, Ronan Le Goffic for the gift of the H7N1-Nluc and the H3N2-Nluc viruses. We thank Nicolas Meunier for his help in statistical analyses. M.B. acknowledges fellowships of the DIM1Health and of the Animal Health Division of INRAE, J.M. acknowledges fellowships of the ANR program and the Animal Health Division of INRAE. B.D. acknowledges support of the ANR-17-CE18-0006-01 program. This work used the platforms of the Grenoble Instruct-ERIC Center (ISBG: UMS 3518 CNRS-CEA-UGA-EMBL) with support from FRISBI (ANR-10-INBS-05-02) and GRAL (ANR-10-LABX-49-01) within the Grenoble Partnership for Structural Biology (PSB). Authors acknowledge the SPR/BLI platform scientific responsible, Jean-Baptiste Reiser Ph.D., for its help and assistance. The funders had no role in study design, data collection and analysis, decision to publish, or preparation of the manuscript.

---

**S1 Figure:** Effect of intracellular VHH expression on H3N2 FluPol activity. Plasmids expressing NP, PA, PB1, PB2 of the H3N2 strain were co-transfected in HEK-293T cells together with the NA-firefly-luciferase reporter plasmid and a plasmid encoding a VHH.

**S2 Figure**: Effect of the VHHs transiently expressed on the replication of WSN(H1N1)-Nluc influenza virus in HEK 293T cells. Data are mean ± s.e.m. n=2 independent transfections with n=3 technical replicates. Matt-Whitney test was used to compare replication in the presence and absence of VHHs at 24 hours post-infection.

**S3 Figure:** Visualization of the green fluorescent protein (GFP) in transduced MDCK clones stably expressing GFP (MDCK-GFP) and VHH16-2A-GFP (MDCK-VHH16).

**S4 Figure:** Visualization of the green fluorescent protein in RK13 clones expressing VHH16-2A-GFP in a doxycycline-inducible manner.

**S5 Figure: A.** Subcellular localization of VHH16-HA in MDCK clones expressing VHH16-HA-2A-GFP in a doxycycline-inducible manner. **B.** Two different MDCK cell clones selected for VHH16-2A-GFP gene expression were incubated (or not) with doxycycline and infected with the reporter influenza virus WSN-Luc. Twenty-four hours post-infection, virus replication was quantified by measurement of the luciferase activity. Data are mean ± s.e.m. n=2 independent transfections with n=3 technical replicates.

**S6 Figure:** Quantification of viral RNAs in infected MDCK-VHH16 cells. Specific primers were used for strand-specific real-time RT-PCR using tagged primers for quantification of the vRNA, cRNA, and mRNA of the segment 5.

## REFERENCES

1. EFSA (European Food Safety Authority), ECDC (European Centre for Disease Prevention and Control), EURL (European Reference Laboratory for Avian Influenza), Adlhoch C, Fusaro A, Gonzales JL, Kuiken T, Marangon S, Niqueux É, Staubach C, Terregino C, Aznar I, Chuzhakina K, Muñoz Guajardo I and Baldinelli F, 2022. Scientific report: Avian influenza overview June–September2022. EFSA Journal 2022;20(10):7597,58pp. 10.2903/j.efsa.2022.7597

2. Shaw M, Palese P. 2013. Orthomyxoviridae, p 1151–1185. In Knipe DM, Howley P (ed), Fields virology, 6th ed, vol 1. Lippincott Williams & Wilkins, Philadelphia, PA.

3. Pflug A, Lukarska M, Resa-Infante P, Reich S, Cusack S. 2017. Structural insights into RNA synthesis by the influenza virus transcription-replication machine. Virus Res. 234:103–117. 10.1016/j.virusres.2017.01.013.

4. Boulo S, Akarsu H, Ruigrok RW, Baudin F. 2007. Nuclear traffic of influenza virus proteins and ribonucleoprotein complexes. Virus Res 124: 12–21. 10.1016/j.virusres.2006.09.013.

5. Deng T, Sharps J, Fodor E, Brownlee GG. 2005. In vitro assembly of PB2 with a PB1-PA dimer supports a new model of assembly of influenza A virus polymerase subunits into a functional trimeric complex. J Virol 79:8669–8674. 10.1128/JVI.79.13.8669-8674.2005.

6. Fodor E, Smith M. 2004. The PA subunit is required for efficient nuclear accumulation of the PB1 subunit of the influenza A virusRNApolymerase complex. J Virol 78:9144–9153. 10.1128/JVI.78.17.9144-9153.2004.

7. Huet S, Avilov SV, Ferbitz L, Daigle N, Cusack S, Ellenberg J. 2010. Nuclear import and assembly of influenza A virus RNA polymerase studied in live cells by fluorescence cross-correlation spectroscopy. J Virol 84:1254–1264. 10.1128/JVI.01533-09.

8. MacDonald LA, Aggarwal S, Bussey KA, Desmet EA, Kim B, Takimoto T. 2012. Molecular interactions and trafficking of influenza A virus polymerase proteins analyzed by specific monoclonal antibodies. Virology 426:51–59. 10.1016/j.virol.2012.01.015.

9. Krischuns T, Lukarska M, Naffakh N, Cusack S. 2021. Influenza Virus RNA-Dependent RNA Polymerase and the Host Transcriptional Apparatus. Annu Rev Biochem. 90:321–348. 10.1146/annurev-biochem-072820-100645.

10. Fodor E, Te Velthuis AJW. 2020. Structure and Function of the Influenza Virus Transcription and Replication Machinery. Cold Spring Harb Perspect Med. 10:a038398. 10.1101/cshperspect.a038398.

11. Tomescu AI, Robb NC, Hengrung N, Fodor E, Kapanidis AN. 2014. Single-molecule FRET reveals a corkscrew RNA structure for the polymerase-bound influenza virus promoter. Proc Natl Acad Sci U S A 111:E3335–42. 10.1073/pnas.1406056111.

12. Blaas D, Patzelt E, Kuechler E. 1982. Identification of the cap binding protein of influenza virus. Nucleic Acids Res 10:4803–4812. 10.1093/nar/10.15.4803.

13. Ulmanen I, Broni BA, Krug RM. 1981. Role of two of the influenza virus core P proteins in recognizing cap 1 structures (m7GpppNm) on RNAs and in initiating viral RNA transcription. Proc Natl Acad Sci U S A 78: 7355–7359. 10.1073/pnas.78.12.7355.

14. Reich S, Guilligay D, Pflug A, Malet H, Berger I, Crépin T, Hart D, Lunardi T, Nanao M, Ruigrok RW, Cusack S. 2014. Structural insight into cap-snatching and RNA synthesis by influenza polymerase. Nature 516:361–366. 10.1038/nature14009.

15. Desmyter A, Farenc C, Mahony J, Spinelli S, Bebeacua C, Blangy S, Veesler D, van Sinderen D, Cambillau C. 2013. Viral infection modulation and neutralization by camelid nanobodies. Proc Natl Acad Sci U S A. 110:E1371–9. 10.1073/pnas.1301336110.

16. Muyldermans S. 2021. Applications of nanobodies. Annu Rev Anim Biosci. 9:401–421. 10.1146/annurev-animal-021419-083831.

17. - Soetens E, Ballegeer M, Saelens X. 2020. An Inside Job: Applications of Intracellular Single Domain Antibodies. Biomolecules. 10:1663. 10.3390/biom10121663

18. Traenkle B, Rothbauer U. 2017. Under the Microscope: Single-Domain Antibodies for Live-Cell Imaging and Super-Resolution Microscopy. Front Immunol. 8:1030. 10.3389/fimmu.2017.01030.

19. Vanlandschoot P, Stortelers C, Beirnaert E, Ibanez LI, Schepens B, Depla E, Saelens X. 2011. Nanobodies(R): new ammunition to battle viruses. Antiviral Res 92:389–407. 10.1016/j.antiviral.2011.09.002.

20. Swale C, Monod A, Tengo L, Labaronne A, Garzoni F, Bourhis JM, Cusack S, Schoehn G, Berger I, Ruigrok RW, Crépin T. 2016. Structural characterization of recombinant IAV polymerase reveals a stable complex between viral PA-PB1 heterodimer and host RanBP5. Sci Rep. 6:24727. 10.1038/srep24727.

21. Nguyen VS, Spinelli S, Desmyter A, Le TT, Kellenberger C, Cascales E, Cambillau C, Roussel A. 2015. Production, crystallization and X-ray diffraction analysis of a complex between a fragment of the TssM T6SS protein and a camelid nanobody Acta Crystallogr F Struct Biol Commun. 71:266–71. 10.1107/S2053230X15000709.

22. Tran V, Moser LA, Poole DS, Mehle A. 2013. Highly sensitive real-time in vivo imaging of an influenza reporter virus reveals dynamics of replication and spread. J Virol. 87:13321–9. 10.1128/JVI.02381-13.

23. Da Costa B, Sausset A, Munier S, Ghounaris A, Naffakh N, Le Goffic R, Delmas B. 2015. Temperature-Sensitive Mutants in the Influenza A Virus RNA Polymerase: Alterations in the PA Linker Reduce Nuclear Targeting of the PB1-PA Dimer and Result in Viral Attenuation. J Virol. 89:6376–90. 10.1128/JVI.00589-15.

24. Morel J, Sedano L, Lejal N, Da Costa B, Batsché E, Muchardt C, Delmas B. 2022. The Influenza Virus RNA-Polymerase and the Host RNA-Polymerase II: RPB4 Is Targeted by a PB2 Domain That Is Involved in Viral Transcription. Viruses. 14:518. 10.3390/v14030518.

25. Nilsson-Payant BE, Sharps J, Hengrung N, Fodor E. 2018. The Surface-Exposed PA51-72-Loop of the Influenza A Virus Polymerase Is Required for Viral Genome Replication. J Virol. 92:e00687–18. 10.1128/JVI.00687-18.

26. Hutchinson EC, Orr OE, Man Liu S, Engelhardt OG, Fodor E. 2011. Characterization of the interaction between the influenza A virus polymerase subunit PB1 and the host nuclear import factor Ran-binding protein 5. J Gen Virol. 92:1859–1869. 10.1099/vir.0.032813-0.

27. Keown JR, Zhu Z, Carrique L, Fan H, Walker AP, Serna Martin I, Pardon E, Steyaert J, Fodor E, Grimes JM. 2022. Mapping inhibitory sites on the RNA polymerase of the 1918 pandemic influenza virus using nanobodies. Nat Commun. 13:251. 10.1038/s41467-021-27950-w.

28. Hanke L, Knockenhauer KE, Brewer RC, van Diest E, Schmidt FI, Schwartz TU, Ploegh HL. 2016. The Antiviral Mechanism of an Influenza A Virus Nucleoprotein-Specific Single-Domain Antibody Fragment. mBio. 7:e01569–16. 10.1128/mBio.01569-16.

29. Ashour J, Schmidt FI, Hanke L, Cragnolini J, Cavallari M, Altenburg A, Brewer R, Ingram J, Shoemaker C, Ploegh HL. 2015. Intracellular expression of camelid single-domain antibodies specific for influenza virus nucleoprotein uncovers distinct features of its nuclear localization. J Virol. 89:2792–800. 10.1128/JVI.02693-14.

30. Kao RY, Yang D, Lau LS, Tsui WH, Hu L, Dai J, Chan MP, Chan CM, Wang P, Zheng BJ, Sun J, Huang JD, Madar J, Chen G, Chen H, Guan Y, Yuen KY. 2010. Identification of influenza A nucleoprotein as an antiviral target. Nat Biotechnol. 28:600–5. 10.1038/nbt.1638

31. Meyer L, Sausset A, Sedano L, Da Costa B, Le Goffic R, Delmas B. 2016. Codon Deletions in the Influenza A Virus PA Gene Generate Temperature-Sensitive Viruses. J Virol. 90:3684–93. 10.1128/JVI.03101-15.

32. Walker AP, Fodor E. 2019. Interplay between Influenza Virus and the Host RNA Polymerase II Transcriptional Machinery. Trends Microbiol. 27:398–407. 10.1016/j.tim.2018.12.013.

33. Zhu Z, Fodor E, Keown JR. 2023. A structural understanding of influenza virus genome replication. Trends Microbiol. 31:308–319. 10.1016/j.tim.2022.09.015.

34. Peacock TP, Sheppard CM, Staller E, Barclay WS. 2019. Host Determinants of Influenza RNA Synthesis. Annu Rev Virol. 6:215–233. 10.1146/annurev-virology-092917-043339.

35. Compton AA, Bruel T, Porrot F, Mallet A, Sachse M, Euvrard M, Liang C, Casartelli N, Schwartz O. 2014. IFITM proteins incorporated into HIV-1 virions impair viral fusion and spread. Cell Host Microbe. 16:736–47. 10.1016/j.chom.2014.11.001.

36. Pardon E, Laeremans T, Triest S, Rasmussen SG, Wohlkönig A, Ruf A, Muyldermans S, Hol WG, Kobilka BK, Steyaert J. 2014. A general protocol for the generation of Nanobodies for structural biology. Nat Protoc. 9:674–93. 10.1038/nprot.2014.039.

37. Meyer L, Sausset A, Sedano L, Da Costa B, Le Goffic R, Delmas B. 2016. Codon Deletions in the Influenza A Virus PA Gene Generate Temperature-Sensitive Viruses. J Virol. 90:3684–93. 10.1128/JVI.03101-15

38. Mettier J, Marc D, Sedano L, Da Costa B, Chevalier C, Le Goffic R. 2021. Study of the host specificity of PB1-F2-associated virulence. Virulence. 12:1647–1660. 10.1080/21505594.2021.1933848

39. Heilmann E, Kimpel J, Geley S, Naschberger A, Urbiola C, Nolden T, von Laer D, Wollmann G. 2019. The Methyltransferase Region of Vesicular Stomatitis Virus L Polymerase Is a Target Site for Functional Intramolecular Insertion. Viruses. 11:989. 10.3390/v11110989.

40. Kawakami E, Watanabe T, Fujii K, Goto H, Watanabe S, Noda T, Kawaoka Y. 2011. Strand-specific real-time RT-PCR for distinguishing influenza vRNA, cRNA, and mRNA. J Virol Methods. 173:1–6. 10.1016/j.jviromet.2010.12.014.

